# Conductance-Based Structural Brain Connectivity in Aging and Dementia

**DOI:** 10.1101/2020.09.15.298331

**Authors:** Aina Frau-Pascual, Jean Augustinak, Divya Varadarajan, Anastasia Yendiki, David H. Salat, Bruce Fischl, Iman Aganj, for the Alzheimer’s Disease Neuroimaging Initiative

**Affiliations:** Athinoula A. Martinos Center for Biomedical Imaging, Massachusetts General Hospital, Harvard Medical School, 149 13th St, Charlestown 02129, MA, USA; Computer Science and Artificial Intelligence Laboratory, Massachusetts Institute of Technology, 32 Vassar St, Cambridge 02139, MA, USA

**Keywords:** brain connectivity, conductance, diffusion MRI, Alzheimer’s disease, aging

## Abstract

**Background:** Structural brain connectivity has been shown to be sensitive to the changes that the brain undergoes during Alzheimer’s disease (AD) progression.

**Methods:** In this work, we used our recently proposed structural connectivity quantification measure derived from diffusion MRI, which accounts for both direct and indirect pathways, to quantify brain connectivity in dementia. We analyzed data from the ADNI-2 and OASIS-3 datasets to derive relevant information for the study of the changes that the brain undergoes in AD. We also compared these datasets to the HCP dataset, as a reference, and eventually validated externally on two cohorts of the EDSD database.

**Results:** Our analysis shows expected trends of mean conductance with respect to age and cognitive scores, significant age prediction values in aging data, and regional effects centered among sub-cortical regions, and cingulate and temporal cortices.

**Discussion:** Results indicate that the conductance measure has prediction potential, especially for age, that age and cognitive scores largely overlap, and that this measure could be used to study effects such as anti-correlation in structural connections.

**Impact statement:** This work presents a methodology and a set of analyses that open new possibilities in the study of healthy and pathological aging. The methodology used here is sensitive to direct and indirect pathways in deriving brain connectivity measures from dMRI, and therefore provides information that many state-of-the-art methods do not account for. As a result, this technique may provide the research community with ways to detect subtle effects of healthy aging and AD.

## 1. Introduction

Brain structural connectivity reflects the physical connections through white-matter (WM) axon bundles between different regions of interest (ROIs), and can be measured by diffusion-weighted magnetic resonance imaging (dMRI). Brain connectivity analysis has been proven to be useful in the study of many conditions, such as the effects of aging on the brain (Damoiseaux, 2017; Wu et al., 2013; Fjell et al., 2016), and in the study of disease. Differences in brain connectivity patterns between healthy and diseased populations are potential indicators of changes in the brain “wiring” due to disease processes. In particular, Alzheimer’s disease (AD) has been found to impact structural connectivity (Rose et al., 2000; Daianu et al., 2013; Prasad et al., 2015). The changes that the brain undergoes with aging and AD can be confounded, thereby contributing to a delay in AD diagnosis. Nonetheless, the spatial and temporal patterns of changes in connectivity are different in healthy aging and in AD. Accurate modeling of structural connectivity may therefore reveal the effects of aging and AD progression in WM degeneration, and help to differentiate the two.

### 1.1. Brain changes in aging and dementia

Healthy aging is associated with a moderate decline of some cognitive abilities. AD dementia causes severe deterioration of similar cognitive domains, but also additional cognitive functions, in such a way that it compromises independent living. The abnormal decline preceding AD, with noticeable alterations in cognition yet short of functional independence, is termed mild cognitive impairment (MCI) (Petersen et al., 1999). Both the gray matter (GM) and the WM undergo changes in volume and integrity in healthy and pathological aging, but the affected regions vary. Healthy aging has been found to be related to decline in frontal and temporal regions of the GM. An age-related volume decline has been localized in the prefrontal cortex (PFC), insula, anterior cingulate gyrus, superior temporal gyrus, inferior parietal lobule, and precuneus (Ghosh et al., 2011), as well as in the striatum, caudate, and medial temporal lobe (hippocampus and adjacent, anatomically related cortex, including entorhinal, perirhinal, and parahippocampal cortices) (Raz and Rodrigue, 2006). Other regions, like occipital cortex, are mostly unaffected by aging. Degeneration in WM has been found to often follow an anterior-posterior gradient of fractional anisotropy (FA) reductions, indicating that frontal connections are especially vulnerable (Toepper, 2017). Small and less myelinated fibers are particularly vulnerable to age-related decline, such as fiber tracts whose myelinization is completed later in life (Salat et al., 2005).

In patients with AD dementia, changes in regional volume are not uniform. A significant volume reduction has been found in the hippocampal formation and the entorhinal cortex bilaterally very early in the disease (Thompson et al., 2004), followed by a degeneration in the PFC (Ghosh et al., 2011). In the WM, signal abnormalities (WMSA) have been found in AD in regions such as rostral frontal, inferior temporal, and inferior parietal WM, with a greater volume of WMSA in AD with respect to healthy aging consistently across different ages. In MCI, frontal and temporal regions have been found to have greater WMSA volume with decreasing time-to-AD-conversion (Lindemer et al., 2017).

### 1.2. Brain connectivity changes in aging and dementia

While enabling the segregation and integration of information processing, brain networks are also responsible for widespread effects resulting from local disease-related disruptions, thereby complicating relationships between pathological processes and clinical phenotypes in AD (Tijms et al., 2013). The disconnection model of AD has long been discussed (de LaCoste and White III, 1993), with cumulative evidence associating plaques and tangles with local synaptic disruptions (Arendt, 2009; Takahashi et al., 2010), as well as linking the cognitive dysfunction in AD to dysconnectivity between highly-interrelated brain regions (Delbeuck et al., 2003; Brier et al., 2014; Matthews et al., 2013). Identifying large-scale brain networks that are vulnerable or resilient in aging and AD (by studying the human *connectome*) can therefore reveal underlying disease propagation patterns in the brain and provide connectivity-based biomarkers (Gomez-Ramirez and Wu, 2014) in prodromal AD. Network-based analysis of brain WM connections through dMRI (Basser and Özarslan, 2014; Goveas et al., 2015; Madden et al., 2012) has been proven promising in revealing the structural basis of cognitive changes in AD, MCI, and aging, and discovery of diagnostically and therapeutically important biomarkers. Structural networks have been used to predict the process of disease spread in AD (Raj et al., 2012, 2015), and to distinguish the groups of cognitively normal (CN), MCI, and AD (Prasad et al., 2015; Frau-Pascual et al., 2019b; Aganj et al., 2014; Shao et al., 2012), as well as AD from vascular dementia (Zarei et al., 2009). Relative to CN controls, AD patients have been shown to have significantly lower integrity of association fiber tracts (Rose et al., 2000), weaker cingulum connectivity (Zhang et al., 2007; Huang et al., 2012; Mielke et al., 2009), and structural brain networks with disruption in their rich club organization (Daianu et al., 2016; Lee et al., 2018), and reduced local efficiency (Reijmer et al., 2013; Lo et al., 2010). The performance in memory and executive functioning of AD patients has been inversely correlated to the path length (Reijmer et al., 2013), and network small-worldness has been shown to predict brain atrophy in MCI (Nir et al., 2015). Structural brain networks are affected even in individuals without dementia with the APOE *t:*4 allele (Liu et al., 2013), a genetic AD risk factor.

Connectivity disruption within a brain network is also occasionally accompanied by hyper-connectivity in a reciprocal network. For instance, increased frontal connectivity may be observed alongside reduced temporal connectivity in AD (Supekar et al., 2008; Wang et al., 2007), and inverse relationship has been reported between frontal activity and occipital activity in aging (Davis et al., 2008). Furthermore, AD has been shown to reduce connectivity in the default mode network (DMN) but intensify it at the early stages in the salience network – a collection of regions active in response to emotionally significant stimuli (Seeley et al., 2007; Uddin, 2016) – whereas behavioral variant fronto-temporal dementia (bvFTD) has been shown to attenuate the salience network connectivity but enhance DMN connectivity (Brier et al., 2012; Zhou et al., 2010). Even so, most existing studies monitor such hyper-connectivity with respect to the progression of dementia, but not with respect to the deterioration of other networks.

### 1.3. Contributions of this work

We have previously introduced a method for inferring structural brain connectivity from dMRI using an electrical conductance model (Frau-Pascual et al., 2019b), which accounts for all possible WM pathways, and is solved globally. This method was shown to produce structural connectivity measures that were more strongly correlated with resting-state functional connectivity and more sensitive to AD-related WM degeneration than standard streamline tractography methods did. In this work, we extend our analysis to demonstrate the impact that this new measure of structural brain connectivity could have in the study of aging and AD dementia. To that end, we investigate the relationship of structural connectivity with age and cognitive and volumetric measures, attempt to predict age and cognitive scores from dMRI data, and identify some anti-correlated connections that might help to study compensation. This paper extends our preliminary conference publications (Frau-Pascual et al., 2019a; Aganj et al., 2020); in particular, we have added more data analysis and experiments.

## 2. Methods

### 2.1. Conductance model

In our previous work (Frau-Pascual et al., 2019b)^1^, we extended the heat equation method proposed by O’Donnell et al. (2002) with a combination of differential circuit laws. Our method assigns to each image voxel a local anisotropic conductivity value *D*, which is the diffusion tensor computed from dMRI (Basser et al., 1994). By solving the partial differential equation (PDE), *−∇ ·* (*D∇φ_i,j_*) = *γ_i,j_,* for a given current configuration *γ_i,j_* between a pair of source (*i*) and sink (*j*) voxels (see below), we find the potential map *φ_i,j_* for that specific configuration. *∇* and *∇·* are the gradient and divergence operators, respectively.

Our algorithm solves the PDE for a 1-ampere current (without loss of generality) between a pair of voxels *i* and *j*: *γ_i,j_* = *δ_i_ − δ_j_*, where *δ_k_*(*x*) := *δ*(*x − x_k_*), with *x_k_* the position of voxel *k* and *δ*(*·*) the Dirac delta. To compute ROI-wise conductance, we distribute the currents among the sets of voxels *I* and *J* (the two ROIs) as: 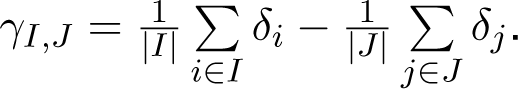

The conductance between two points is then computed from Ohm’s law as the ratio of the current to the potential difference. In our case, the potential difference between two voxels (or ROIs) i and j is _i;j(x_i_) - Ø_i;j_(x_j_). The conductance is therefore computed, for ROI-wise connectivity, as: 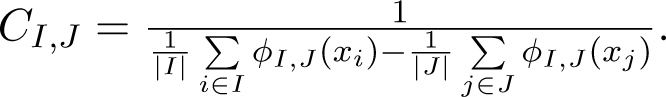

The conductance among all *N* regions can be computed efficiently in O(*N*) using the superposition principle (Frau-Pascual et al., 2019b). High conductance (i.e. low resistance) between two points represents a high degree of connectivity in our model. Since the ROIs are all at least weakly connected, these maps can then be thresholded.

### 2.2. Brain connectivity matrix generation

With the conductance method, we model and quantify diffusion data in a non-conventional way. As mentioned in Section 2.1, a 1-ampere current is split across voxels and the PDE is solved once per ROI to compute a conductance measure between each pair of ROIs using the superposition principle. This eventually results in a connectivity matrix per subject that reflects the ease with which this small current goes from one region to another, following the diffusion tensors. This measure also embeds geometrical information, such as volumes (number of voxels in each ROI) and distances between ROIs (as implied by the spatial derivatives in the PDE).

### 2.3. Study of conductance matrices

In this study, we considered the relationship between the mean conductance and other variables, such as age and cognitive scores (and cortical/subcortical volumes in the Supplementary Materials). The cognitive scores considered here were the Clinical Dementia Rating (CDR) scale (Morris, 1991) and the Mini–Mental State Examination (MMSE) score (Pangman et al., 2000), explained in more detail in Section 3.1. We fit a linear function to our data points and report the correlation (*r*) and significance (*p*) values of the fit, revealing whether conductance was significantly correlated with these variables.

We also attempted to *predict* variables such as age and cognitive scores via linear regression. We initially discarded outliers in every cohort, while considering an outlier a subject with mean conductance higher or lower than the average by two standard deviations. We report *r* and *p* values of the correlation between the predicted and the true variable when fitting with a cohort and predicting with a different cohort. The *p*-values below 0.05 were considered significant when *r* was positive (as a negative *r* would have indicated the opposite of the desired effect). We also report the Bonferroni corrected *p*-values (*p_b_*), i.e. the original *p*-values multiplied by the number of comparisons. Furthermore, we tried fitting and predicting within the same cohort (which involved fewer comparisons), using cross-validation with 20 folds.

We then measured the correlation of the conductance values between every pair of regions with age and cognitive scores (CDR and MMSE). For each pair of regions, we corrected the *p*-value for multiple comparisons using Bonferroni correction. The results would reveal which regions correlate more strongly with age and/or cognition.

We further used a general linear model to regress out the effects of age and sex before correlating the conductance with CDR or MMSE. This would disentangle the overlapping contributions of age/sex and cognition, and reveal the residual effects of CDR and MMSE unexplained by age/sex.

### 2.4. Identification of anti-correlated connections

Next, we attempted to identify negative (cross-subject) interrelationships among brain connections. As opposed to focusing only on the relationship between connectivity and the clinical data, we identified pairs of connections that are significantly negatively correlated with each other, and validated them on external datasets. Such a connection-wise correlation approach might help to reveal pathways that are potentially compensatory and define the resilience mechanism of brain networks against AD.

We first vectorized the lower triangular part of each *N × N* connectivity matrix to a vector of length *N* (*N −* 1)*/*2, and reduced this vector to keep *M* cortico-cortical and cortico-subcortical connections. Next, we computed the cross-subject linear correlation coefficient between each pair of connections, resulting in two symmetric *M ×M* connection-wise matrices of correlations, *R*, and *p*-values, *P*. We then kept only the elements of *R* with a correlation value smaller than a negative threshold, e.g. *τ* = *−*0.1, as *R^−^*= *{*(*i, j*)*|R_i,j_< τ}*. From that set, we considered the connection pairs whose *p*-values survived a cutoff threshold, namely *α* = 0.05, as *L* = *{*(*i, j*) *∈ R^−^|P_i_^∗^_,j_< α}*. *P^∗^*was the set of *p*-values corrected for multiple comparisons among the elements of *R^−^*with the Holm-Bonferroni method. We regarded the surviving set *L* as the pairs of connections with significant cross-subject anti-correlation. We kept either the entire *L*, or a most significant subset of it.

Next, to *externally* test if the surviving set *L* was anti-correlated, we computed *R_test_* and *P_test_* for the connection pairs in *L* in a different population, and verified both *R_test_ <* 0 and *P_t_^∗^_est_ < α* for that set, with *P_t_^∗^_est_* being *P_test_* corrected for multiple comparisons among the pairs in *L*. We also tested the hypothesis that the surviving pairs of connections were left-right symmetric; i.e., whether a significant anti-correlation was also a significant anti-correlation in the mirrored hemisphere.

Lastly, we correlated the identified connections with cognitive performance measures.

### 2.5. MR data processing

The common pipeline for brain connectivity computation is: segmentation of brain ROIs, quantification of brain connections from dMRI, and aggregation of connectivity values in a matrix. The constructed brain connectivity matrix describes how strongly different regions are connected to each other according to the dMRI acquisition of WM connections. We processed the MRI data similarly for all datasets.

#### 2.5.1. Structural MRI processing

We performed tissue segmentation and parcellation of the cortex into ROIs using FreeSurfer^2^ (Fischl, 2012). The parcellation used in this work was the Desikan-Killiany atlas (Desikan et al., 2006), which has 86 regions, among which 68 were cortical and 18 were subcortical or brainstem. The atlas used here has a moderate number of ROIs, which helps to preserve statistical power after Bonferroni correction. For stability and replicability, we decided to use the segmentation results already provided by the database staff (except for EDSD), rather than to reprocess all the structural images with the latest version of FreeSurfer.

#### 2.5.2. Diffusion MRI processing

Diffusion preprocessing was performed using the FSL software^3^ (Jenkinson et al., 2012) and included BET for brain extraction and EDDY for eddy current and subject motion correction.^4^ From the preprocessed dMR images, we reconstructed the diffusion tensors using the Diffusion Tensor Imaging (DTI) (Basser et al., 1994) reconstruction module of DSI Studio^5^, which we then used as input to our conductance computation algorithm.

## 3. Results

### 3.1. Analysis of AD population

We evaluated how our conductance method could help in evaluating the AD stage. For this, we used two publicly available datasets that included subjects across the AD dementia spectrum (see Figure 1): the second phase of Alzheimer’s Disease Neuroimaging Initiative (ADNI-2)^6^ (Jack et al., 2008; Beckett et al., 2015), and the third release in the Open Access Series of Imaging Studies (OASIS-3) (Fotenos et al., 2005), which is a longitudinal neuroimaging, clinical, and cognitive dataset for normal aging and AD. To avoid data heterogeneity, we divided the OASIS-3 dataset into 4 cohorts, each of which included more than 100 subjects with similar description in the “Scans” field of the data sheet. These two datasets, which we used for training and internal validation, enabled us to compare structural brain connectivity in different stages of the disease and correlate neuroimaging data to clinical cognitive scores. We also compared these two datasets to 100 subjects of the publicly available Human Connectome Project (HCP) (Van Essen et al., 2013), which contains data of younger healthy subjects and provides a reference, helping us to interpret our results in the targeted population. Lastly, we evaluated our predictive models on held-out subjects from the European DTI Study in Dementia (EDSD) (Brueggen et al., 2017), including two cohorts imaged in the cities of Freiburg and Rostock, which were chosen due to their within-cohort consistency of image acquisition parameters.

**Figure 1:**
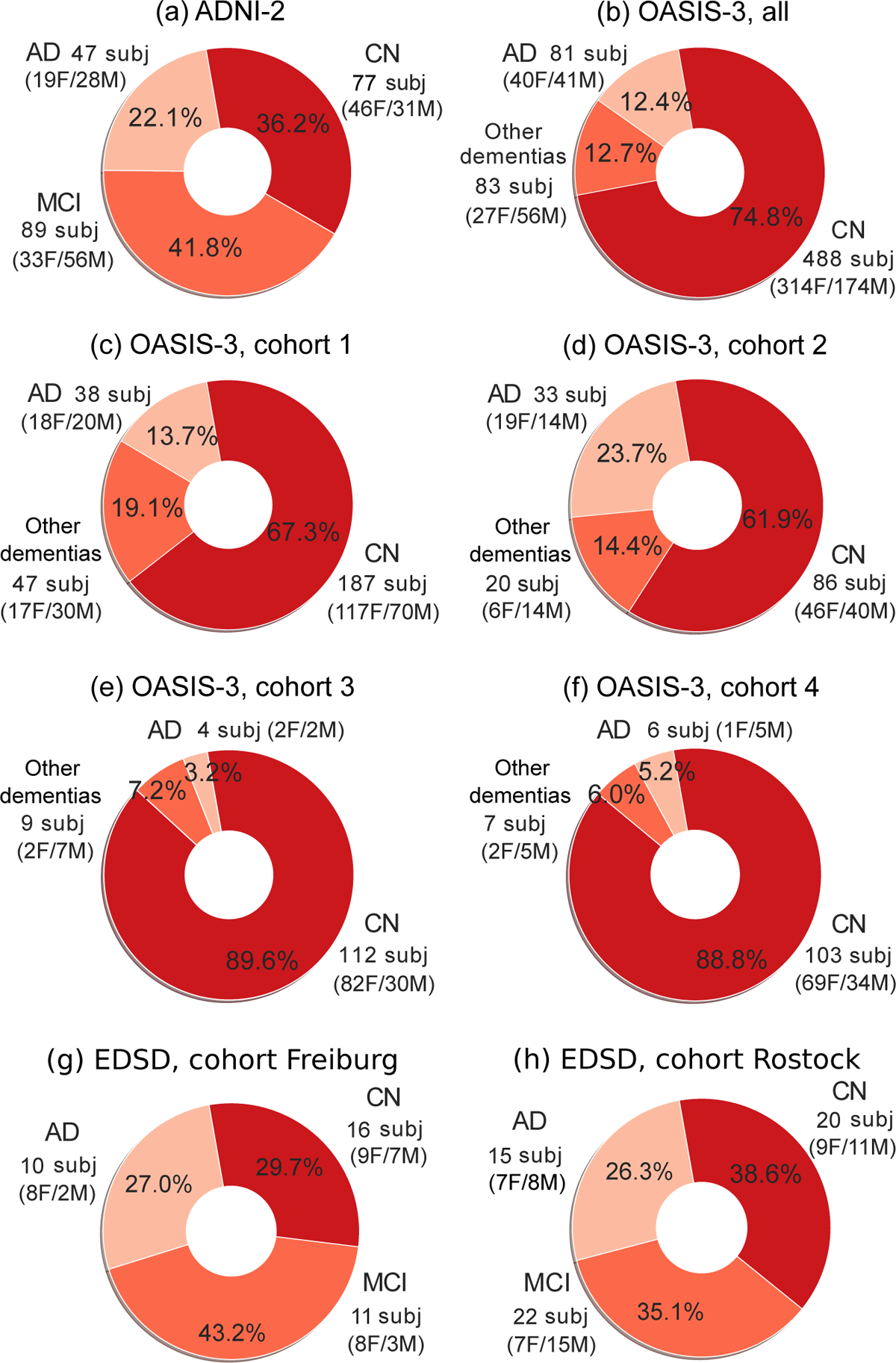
Demographics of the AD datasets used here. (a) ADNI-2 cohort of 213 subjects (77 CN, 89 MCI, 47 AD), (b) OASIS-3 group of 652 subjects, consisting of 4 cohorts, each of which had more than 100 subjects with similar description in the “Scans” field of the data sheet: (c) 272-subject cohort (187 CN, 38 AD, 47 other dementias), (d) 139-subject cohort (86 CN, 33 AD, 20 other dementias), (e) 125-subject cohort (112 CN, 4 AD, 9 other dementias), and (f) 116-subject cohort (103 CN, 6 AD, 7 other dementias). Other dementias included vascular dementia, or AD dementia with depression or additional symptoms (refer to Figure 2). (g) EDSD Freiburg cohort (16 CN, 11 MCI, 10 AD). (h) EDSD Rostock-3T cohort (20 CN, 22 MCI, 15 AD).

The data used in this work, as shown in Table 1, is quite heterogeneous in terms of acquisition, even though all MR images were acquired at 3T. Nevertheless, the conductance method uses only the simple diffusion tensor (as opposed to a higher-order) model, whose performance has been shown to stabilize after the asymptotic limit of 30 gradient orientations (Jones, 2004). As seen in

Table 1, the number of gradient orientations is at least 23 for all our subjects and we expect the derived diffusion tensors to be robust enough for a fair comparison.

Demographic and clinical data from these populations were also available: age, sex, diagnosis, cerebral cortical and subcortical volumes, and cognitive scores such as the CDR scale (Morris, 1991) and the MMSE score (Pangman et al., 2000). CDR measures from 0 to 3 the cognitive capabilities of each subject, with 0 being CN and a higher number reflecting higher cognitive impairment. MMSE rates cognitive capabilities from 0 to 30, with 30 being CN and a lower value reflecting higher cognitive impairment. Figure 2 shows the relationship between these scores and the subject diagnosis in ADNI-2 and OASIS-3. It is to be noted that the ratings differ across diagnoses and datasets. ADNI-2 rates people with AD diagnosis with CDR scales of 0.5 and 1, MCI with 0.5, and CN with 0, but the MMSE scores are overlapping for the three diagnoses. OASIS-3 rates are variable, with MMSE values overlapping across diagnoses and CDR scores. Therefore, the ratings across the datasets are slightly different, and the cognitive scores of MMSE and CDR are used differently.

**Figure 2:**
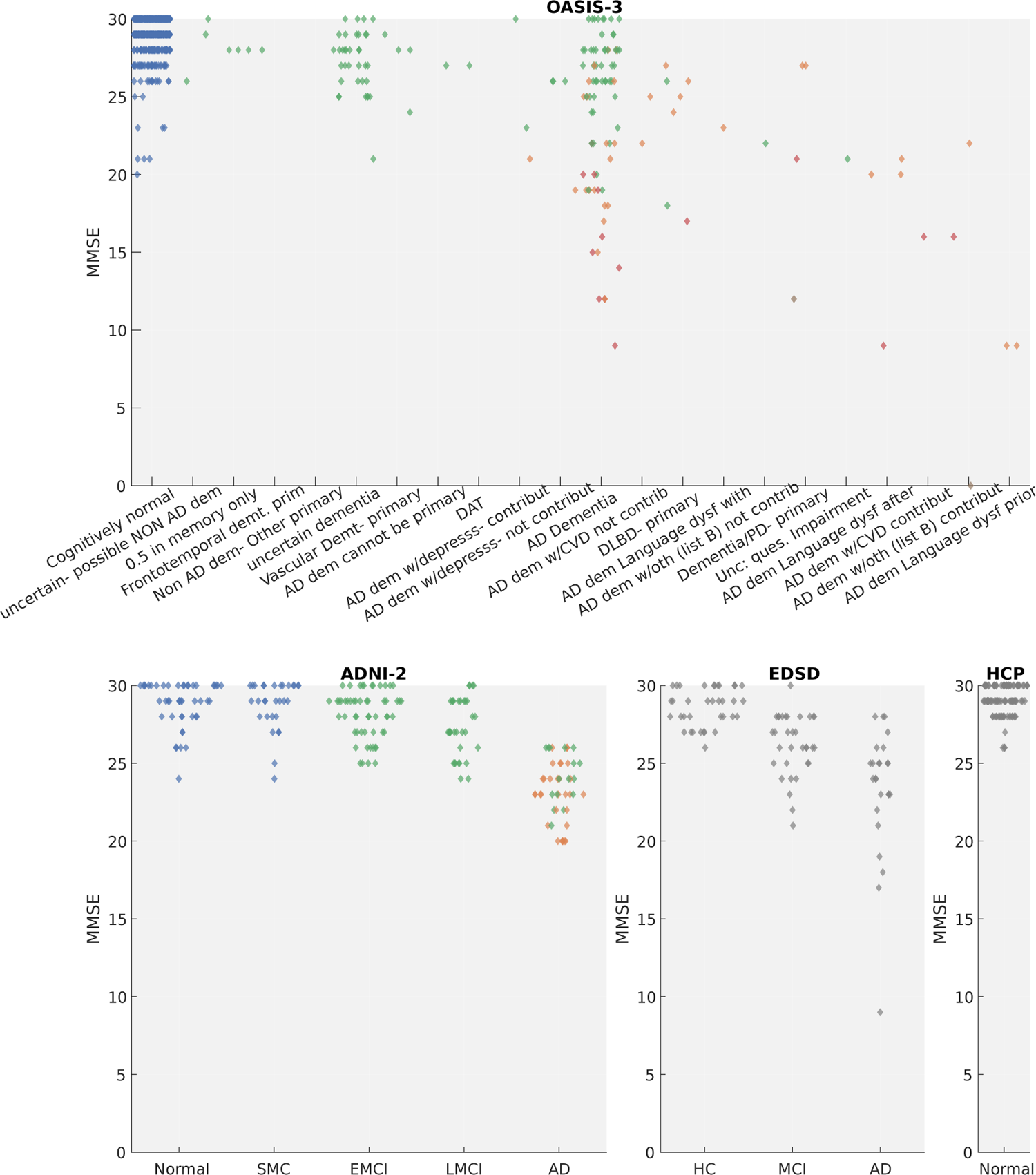
Distribution of the MMSE score for each diagnostic category in each dementia database, color-coded by the CDR values of 0 (blue), 0.5 (green), 1 (orange), 2 (red) and 3 (brown) (refer to Figure 3, right, for the colormap). Diagnostic labels are quoted from the databases. CDR was not available for HCP and EDSD.

### 3.2. Correlation of conductance values with clinical data

We computed the correlation between mean conductance and other variables, such as age, CDR and MMSE (shown in Figure 3), and cortical/subcortical volumes (provided in the Supplementary Materials). Mean conductance consistently exhibited a decreasing trend (negative *r*) with respect to age and CDR and mostly an increasing trend (positive *r*) with respect to MMSE.

**Figure 3:**
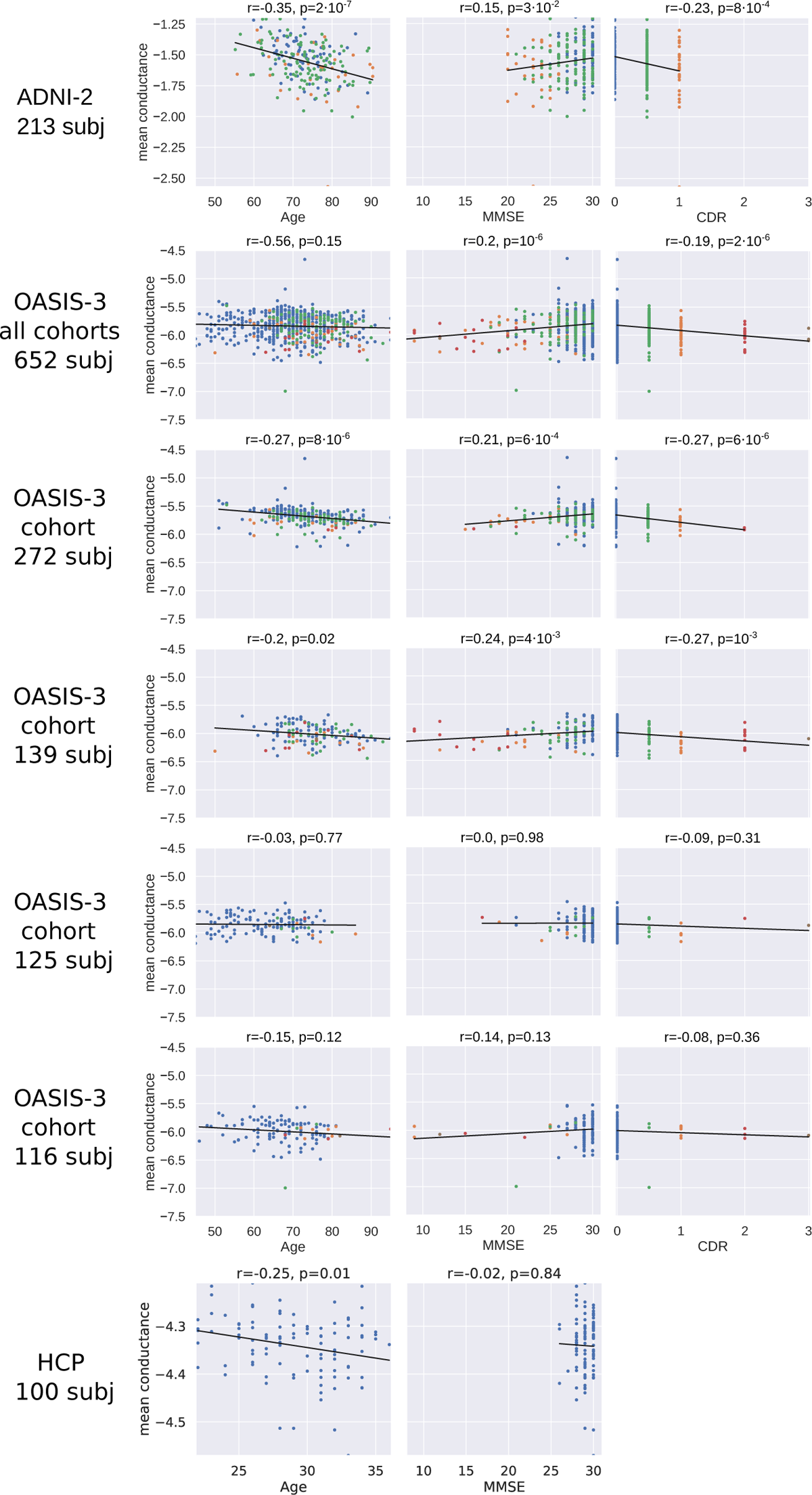
Correlation of mean conductance with age and cognitive scores of CDR and MMSE, color-coded with respect to the CDR: 0 (blue), 0.5 (green), 1 (orange), 2 (red) and 3 (brown). CDR was not available for HCP.

### 3.3. Predictive value of conductance matrices

We then assessed the predictive value of our conductance matrices. We used linear regression to fit on one cohort and predict from a different one, or fit and predict on the same cohort using cross-validation (see Section 2.3). We first discarded outliers – subjects with mean conductance higher or lower than the average by two standard deviations – in every cohort. We removed 3 subjects of the ADNI-2 cohort, 3, 6, 2, and 6 subjects of the different OASIS-3 cohorts, and no subject from the HCP cohort.

In tables 2 and 3, we show the *r* and *p* values when fitting and predicting age with the same cohort (within-cohort prediction), and when fitting age with a cohort and predicting with a different cohort (cross-prediction), respectively. Table 3 shows significant correlation between predicted and true values of age across ADNI-2 and OASIS-3, but not when fitting using HCP. When we combined all OASIS-3 data, we got values of *r* = 0.212, *p* = 7 *·* 10*^−^*^8^, *p_b_* = 2 *·* 10*^−^*^7^ for within-cohort prediction, and values of *r* = 0.383, *p* = 7 *·* 10*^−^*^9^, *p_b_* = 4 *·* 10*^−^*^8^ when training on OASIS-3 and testing on ADNI-2, and *r* = 0.265, *p* = 10*^−^*^11^, *p_b_* = 6 *·* 10*^−^*^11^ when training on ADNI-2 and testing on OASIS-3, and a negative *r* when training with OASIS-3 and testing on HCP and vice versa. In this comparison (with all of OASIS-3 in a single cohort), we Bonferroni-corrected (*p_b_*) with a factor 3 in the within-cohort prediction and a factor 6 in the cross-prediction case.

**Table 1:** dMRI acquisition parameters for the datasets used.

| Dataset | N. Gradient Orientations | b-value (sec/mm<sup>2</sup>) | Voxel Size (mm<sup>3</sup>) | Field Strength (Tesla) |
| --- | --- | --- | --- | --- |
| ADNI-2 | 41 | 1000 | 1.37×1.37×.7 | 3 |
| OASIS-3 | 23 – 64 | ≤1000 – ≤1400 | 1.22×1.22 – 2.5×2.5, ×2 – ×4 | 3 |
| HCP | 270 | 1000 + 2000 + 3000 | 1.25×1.25×1.25 | 3 |
| EDSD - Freiburg | 61 | 1000 | 2×2×2 | 3 |
| EDSD - Rostock | 60 | 1000 | 0.98×0.98×2.4 | 3 |

**Table 2:** Prediction of **age** within ADNI-2, OASIS-3, and HCP. *pb* stands for Bonferroni-corrected *p*-value (for 6 comparisons). *p*-values under 0.05 with a positive *r* are highlighted.

| ADNI-2 |  | OASIS-3 |  |  | HCP |
| --- | --- | --- | --- | --- | --- |
| 213 sub | 272 sub | 139 sub | 125 sub | 116 sub | 100 sub |
| $r=0.33$,<br>$p=8 \cdot 10^{-7}$,<br>$p_b=5 \cdot 10^{-6}$ | $r=0.14$,<br>$p=0.02$,<br>$p_b=0.14$ | $r=0.34$,<br>$p=6 \cdot 10^{-5}$,<br>$p_b=4 \cdot 10^{-4}$ | $r=0.56$,<br>$p=10^{-11}$,<br>$p_b=7 \cdot 10^{-11}$ | $r=0.33$,<br>$p=5 \cdot 10^{-4}$,<br>$p_b=3 \cdot 10^{-3}$ | $r=-0.06$,<br>$p=0.55$,<br>$p_b=1$ |

**Table 3:** Prediction of **age** in ADNI-2, OASIS-3, and HCP. *pb* stands for Bonferroni-corrected *p*-value (for 30 comparisons). *p*-values under 0.05 with a positive *r* are highlighted.

| Predict | ADNI-2 |  | OASIS-3 |  |  | HCP |
| --- | --- | --- | --- | --- | --- | --- |
| Fit | 213 sub | 272 sub | 139 sub | 125 sub | 116 sub | 100 sub |
| ADNI-2<br>213 sub | | $r=0.25$,<br>$p=4 \cdot 10^{-5}$,<br>$p_b=10^{-3}$ | $r=0.22$,<br>$p=0.01$,<br>$p_b=0.33$ | $r=0.29$,<br>$p=10^{-3}$,<br>$p_b=0.03$ | $r=0.23$,<br>$p=0.02$,<br>$p_b=0.51$ | $r=0.07$,<br>$p=0.51$,<br>$p_b=1$ |
| OASIS-3<br>272 sub | $r=0.34$,<br>$p=3 \cdot 10^{-7}$,<br>$p_b=10^{-5}$ | | $r=0.25$,<br>$p=4 \cdot 10^{-3}$,<br>$p_b=0.11$ | $r=0.26$,<br>$p=4 \cdot 10^{-3}$,<br>$p_b=0.11$ | $r=0.29$,<br>$p=3 \cdot 10^{-3}$,<br>$p_b=0.08$ | $r=-0.18$,<br>$p=0.08$,<br>$p_b=1$ |
| OASIS-3<br>139 sub | $r=0.36$,<br>$p=6 \cdot 10^{-8}$,<br>$p_b=2 \cdot 10^{-6}$ | $r=0.26$,<br>$p=2 \cdot 10^{-5}$,<br>$p_b=5 \cdot 10^{-4}$ | | $r=0.29$,<br>$p=10^{-3}$,<br>$p_b=0.04$ | $r=0.26$,<br>$p=7 \cdot 10^{-3}$,<br>$p_b=0.2$ | $r=0.12$,<br>$p=0.24$,<br>$p_b=1$ |
| OASIS-3<br>125 sub | $r=0.37$,<br>$p=2 \cdot 10^{-8}$,<br>$p_b=6 \cdot 10^{-7}$ | $r=0.2$,<br>$p=10^{-3}$,<br>$p_b=0.03$ | $r=0.33$,<br>$p=9 \cdot 10^{-5}$,<br>$p_b=3 \cdot 10^{-3}$ | | $r=0.35$,<br>$p=2 \cdot 10^{-4}$,<br>$p_b=5 \cdot 10^{-3}$ | $r=-0.02$,<br>$p=0.8$,<br>$p_b=1$ |
| OASIS-3<br>116 sub | $r=0.49$,<br>$p=2 \cdot 10^{-14}$,<br>$p_b=6 \cdot 10^{-13}$ | $r=0.41$,<br>$p=3 \cdot 10^{-12}$,<br>$p_b=8 \cdot 10^{-11}$ | $r=0.31$,<br>$p=2 \cdot 10^{-4}$,<br>$p_b=7 \cdot 10^{-3}$ | $r=0.34$,<br>$p=10^{-4}$,<br>$p_b=3 \cdot 10^{-3}$ | | $r=0.28$,<br>$p=4 \cdot 10^{-3}$,<br>$p_b=0.12$ |
| HCP<br>100 sub | $r=-0.33$,<br>$p=5 \cdot 10^{-7}$,<br>$p_b=10^{-5}$ | $r=-0.16$,<br>$p=8 \cdot 10^{-3}$,<br>$p_b=0.25$ | $r=-0.1$,<br>$p=0.24$,<br>$p_b=1$ | $r=-0.19$,<br>$p=0.03$,<br>$p_b=0.93$ | $r=-0.3$,<br>$p=10^{-3}$,<br>$p_b=0.04$ | |

Tables 4, 5, 6, and 7 show results when similarly predicting the MMSE and CDR cognitive scores. Prediction in these cases yielded less significant *p*-values, both in within- and cross-cohort predictions. When we put all the OASIS-3 data together, we got significant values for within-cohort prediction: *r* = 0.132, *p* = 9 *·* 10*^−^*^4^, and *p_b_* = 2 *·* 10*^−^*^3^ for CDR, *r* = 0.091, *p* = 0.02, and *p_b_* = 0.06 for MMSE. For cross-prediction with ADNI-2: *r* = 0.321, *p* = 2 *·* 10*^−^*^6^, *p_b_* = 4 *·* 10*^−^*^6^ when training on OASIS-3 and testing on ADNI-2, and *r* = 0.054, *p* = 0.17, *p_b_* = 0.34 when training on ADNI-2 and testing on OASIS-3, when using CDR; *r* = 0.317, *p* = 2 *·* 10*^−^*^6^, *p_b_* = 10*^−^*^5^ when training on OASIS-3 and testing on ADNI-2, and *r* = 0.01, *p* = 0.83, *p_b_* = 1 when training on ADNI-2 and testing on OASIS-3, when using MMSE. *p*-values were significant only in one direction: when we trained with all OASIS-3 data together. CDR was not available for the HCP subjects, and a factor 2 was used in the Bonferroni correction in both the within-cohort prediction and cross-prediction. In the case of MMSE, we had the values for HCP too, and when comparing with all of OASIS-3 in a single cohort, we Bonferroni-corrected (*p_b_*) with a factor 3 in the within-cohort prediction and a factor 6 in the cross-prediction case. We observed no significant values in all cases with positive *r* (as negative *r* would not indicate prediction).

**Table 4:**
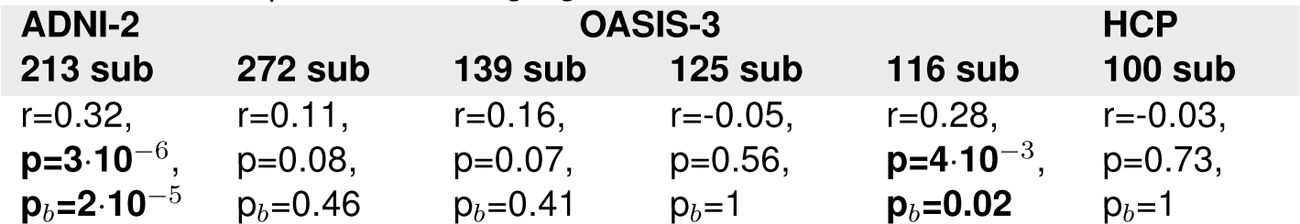
Prediction of **MMSE** within ADNI-2, OASIS-3, and HCP. *pb* stands for Bonferroni-corrected *p*-value (for 6 comparisons). *p*-values under 0.05 with a positive *r* are highlighted.

**Table 5:**
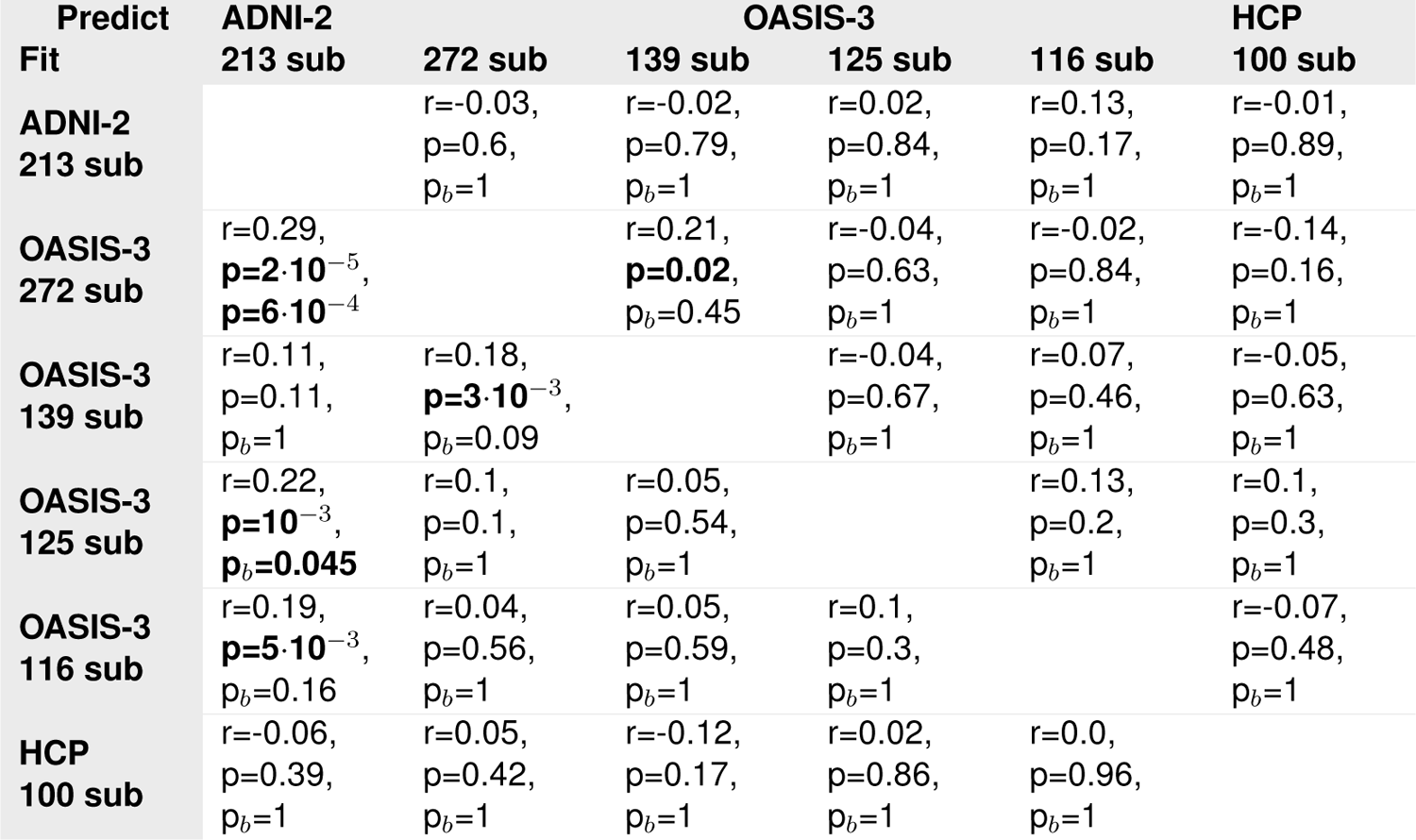
Prediction of **MMSE** in ADNI-2, OASIS-3, and HCP. *pb* stands for Bonferroni-corrected *p*-value (for 30 comparisons). *p*-values under 0.05 with a positive *r* are highlighted.

**Table 6:**
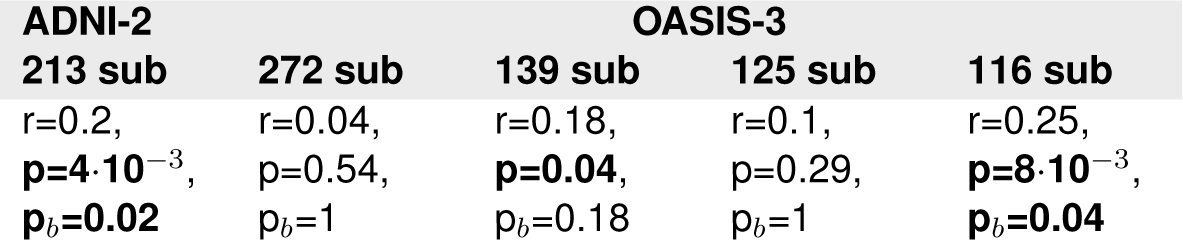
Prediction of **CDR** within ADNI-2 and OASIS-3. *pb* stands for Bonferroni-corrected *p*-value (for 5 comparisons). *p*-values under 0.05 with a positive *r* are highlighted.

| ADNI-2 |  | OASIS-3 |  |  |
| --- | --- | --- | --- | --- |
| 213 sub | 272 sub | 139 sub | 125 sub | 116 sub |
| $r=0.2$,<br>**$p=4 \cdot 10^{-3}$**,<br>**$p_b=0.02$** | $r=0.04$,<br>$p=0.54$,<br>$p_b=1$ | $r=0.18$,<br>**$p=0.04$**,<br>$p_b=0.18$ | $r=0.1$,<br>$p=0.29$,<br>$p_b=1$ | $r=0.25$,<br>**$p=8 \cdot 10^{-3}$**,<br>**$p_b=0.04$** |

**Table 7:** Prediction of **CDR** in ADNI-2 and OASIS-3. *pb* stands for Bonferroni-corrected *p*-value (for 20 comparisons). *p*-values under 0.05 with a positive *r* are highlighted.

| Predict | ADNI-2 |  | OASIS-3 |  |  |
| --- | --- | --- | --- | --- | --- |
| Fit | 213 sub | 272 sub | 139 sub | 125 sub | 116 sub |
| **ADNI-2**<br>**213 sub** | | $r=-0.01$,<br>$p=0.83$,<br>$p_b=1$ | $r=0.03$,<br>$p=0.75$,<br>$p_b=1$ | $r=0.15$,<br>$p=0.1$,<br>$p_b=1$ | $r=0.02$,<br>$p=0.85$,<br>$p_b=1$ |
| **OASIS-3**<br>**272 sub** | $r=0.25$,<br>**$p=3 \cdot 10^{-4}$**,<br>**$p_b=5 \cdot 10^{-3}$** | | $r=0.04$,<br>$p=0.62$,<br>$p_b=1$ | $r=0.08$,<br>$p=0.41$,<br>$p_b=0.82$ | $r=0.1$,<br>$p=0.32$,<br>$p_b=1$ |
| **OASIS-3**<br>**139 sub** | $r=0.2$,<br>**$p=4 \cdot 10^{-3}$**,<br>$p_b=0.08$ | $r=0.18$,<br>**$p=4 \cdot 10^{-3}$**,<br>$p_b=0.08$ | | $r=0.07$,<br>$p=0.45$,<br>$p_b=1$ | $r=0.11$,<br>$p=0.28$,<br>$p_b=1$ |
| **OASIS-3**<br>**125 sub** | $r=0.28$,<br>**$p=4 \cdot 10^{-5}$**,<br>**$p_b=8 \cdot 10^{-4}$** | $r=0.27$,<br>**$p=6 \cdot 10^{-6}$**,<br>**$p_b=10^{-4}$** | $r=0.08$,<br>$p=0.36$,<br>$p_b=1$ | | $r=0.16$,<br>$p=0.09$,<br>$p_b=1$ |
| **OASIS-3**<br>**116 sub** | $r=0.33$,<br>**$p=9 \cdot 10^{-7}$**,<br>**$p_b=2 \cdot 10^{-5}$** | $r=0.19$,<br>**$p=10^{-3}$**,<br>**$p_b=0.03$** | $r=0.14$,<br>$p=0.1$,<br>$p_b=1$ | $r=0.13$,<br>$p=0.16$,<br>$p_b=1$ | |

### 3.4. Region-specific conductance results

We then correlated the elements of the conductance matrix individually with age, the MMSE score, and CDR, while correcting for multiple comparisons by multiplying the *p*-values by the number of connections. Figure 4 shows many of the connections being significant (i.e., *p_b_ <* 0.05) for OASIS-3 and ADNI-2. In the case of age, 1342 and 1981 connections out of 3655 (see Table 8) had *p*-values under 0.05, respectively. Regarding the cognitive scores: for CDR, 683 and 807 connections out of 3655 and for MMSE, 653 and 474 connections out of 3655 had *p*-values under 0.05, in OASIS-3 and ADNI-2, respectively. It is worth noting that in general we found more significant values in OASIS-3, possibly due to the larger sample size. In OASIS-3, age correlated with connectivity with higher significance among all subcortical regions, especially thalamus and hippocampus, bilaterally, and with regions in the cortex, mostly transverse temporal, cingulate regions, insula, and precuneus cortex, bilaterally. In ADNI-2, age also correlated with higher significance in connections involving thalamus and hippocampus, bilaterally, and with regions in the cortex such as middle and superior temporal, lateral orbitofrontal, posterior cingulate, entorhinal, fusiform, and some other regions in occipital and parietal cortices. In OASIS-3, connections found to most significantly correlate with cognitive scores were similar to those correlating with age the most, namely connecting hippocampus, amygdala, insula, and transverse temporal. In ADNI-2, the correlation with age and cognitive scores were different: age was significantly correlated to connections between many regions, whereas the correlation of cognitive scores was significant with connections from hippocampus and amygdala to temporal and cingulum lobes, as well as orbitofrontal.

**Table 8:** Number of significant (surviving) connections (out of 3655).

|  | OASIS-3 | ADNI-2 |
| --- | --- | --- |
| Age | 1342 | 1981 |
| MMSE | 653 | 474 |
| CDR | 683 | 807 |
| MMSE after regressing out age and sex | 282 | 91 |
| CDR after regressing out age and sex | 417 | 518 |

**Figure 4:**
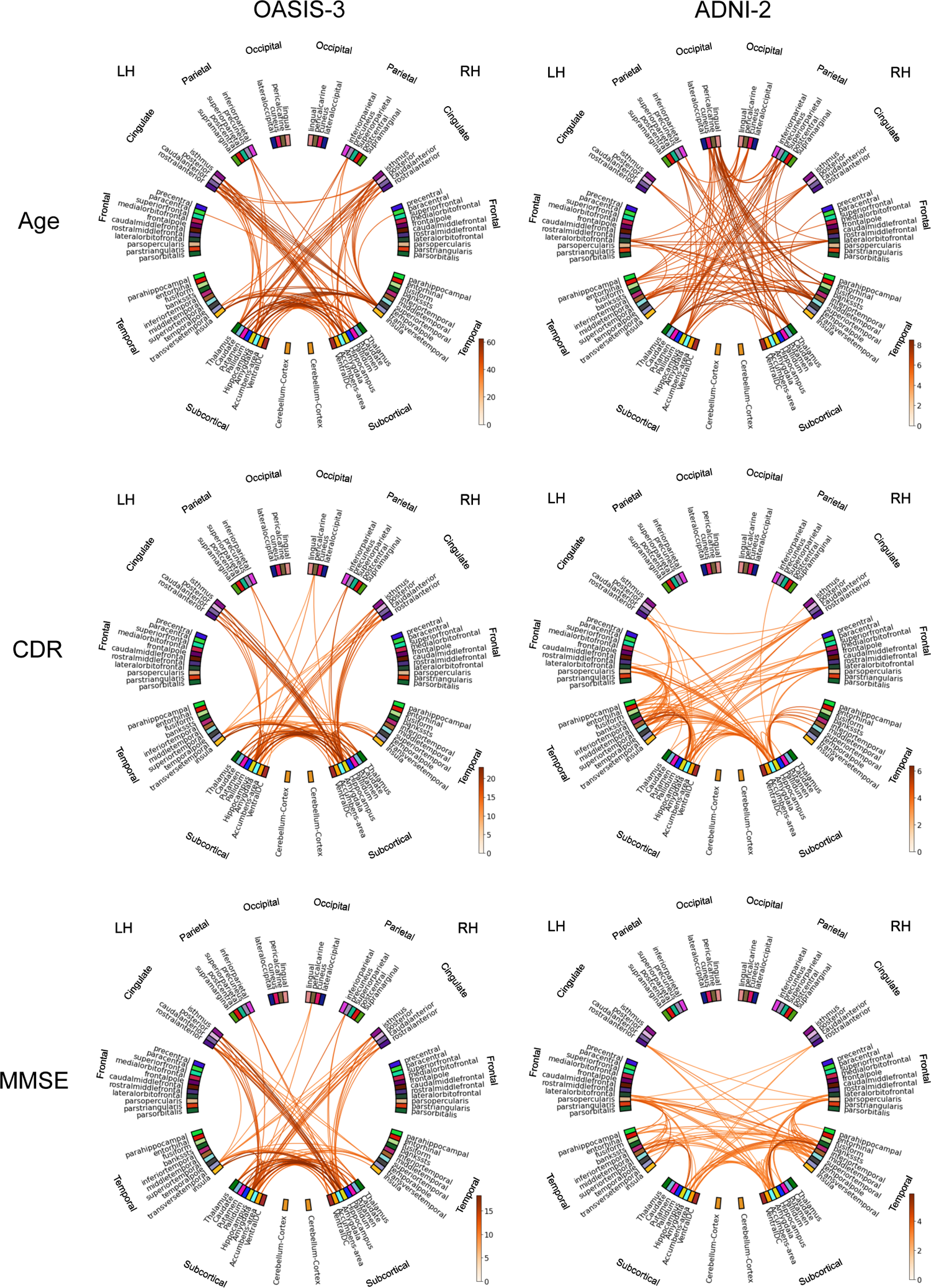
*Sig*-values for the correlation of conductance with age, CDR, and MMSE. We depict the negative logarithm of the Bonferroni-corrected *p*-value (*sig* = *−* log_10_(*pb*)), and consider significant values above 1.3 (i.e., *pb <* 0.05). In this figure, all OASIS-3 cohorts were used together.

We then regressed out the age and sex effects before correlating the conductance with the CDR and MMSE scores. By doing so, the general significance levels decreased. The number of significant connections out of 3655 went down, in OASIS-3, from 683 to 417 for CDR and from 653 to 282 for MMSE and, in ADNI-2, from 807 to 518 for CDR and from 474 to 91 for MMSE. The connectivity plots for this experiment are depicted in Figure 5, showing the most significant connections to be similar to those in Figure 4, but with a lower significance level in general.

**Figure 5:**
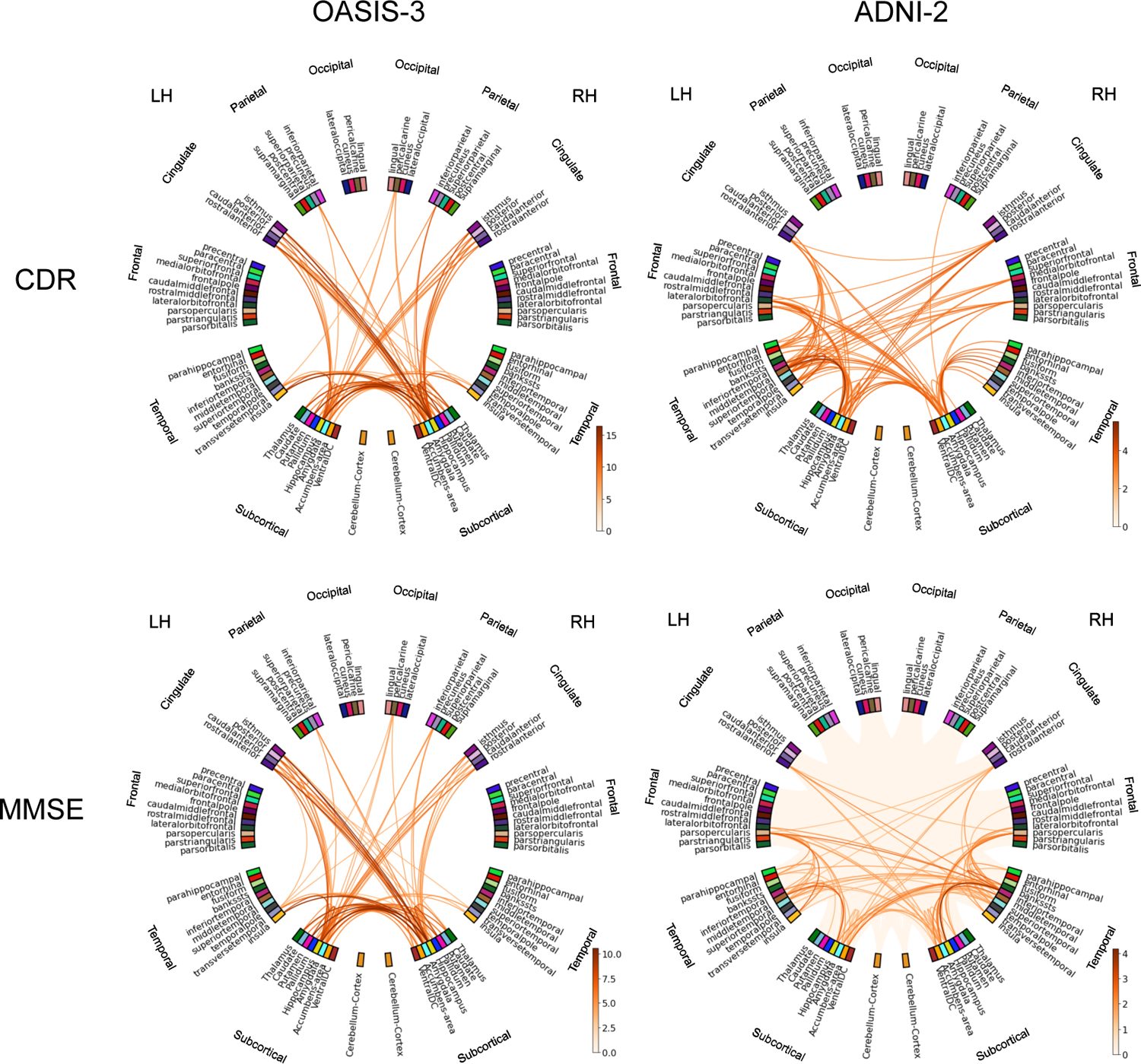
*Sig*-values for the correlation of conductance with CDR and MMSE, once the age and sex effects have been removed. We show the negative logarithm of the Bonferroni-corrected *p*-value (*sig* = − log_10_(*pb*)), and consider significant values above 1.3 (i.e., *pb <* 0.05). In this figure, all OASIS-3 cohorts were used together.

### 3.5. External validation on held-out data

For a true external validation on unseen data (without fine-tuning the pipeline), we applied the predictive models that we trained using ADNI-2, OASIS-3, and HCP on the two held-out EDSD cohorts. We removed 4 subjects from each of the Freiburg and Rostock cohorts due to failure in structural image processing. The results of the prediction of age and MMSE are reported in Table 9. CDR was not available for EDSD.

**Table 9:** Prediction of **age** and **MMSE** in the held-out EDSD database. *pb* stands for Bonferroni-corrected *p*-value (for 14 comparisons for each variable). *p*-values under 0.05 with a positive *r* are highlighted.

| Predict | EDSD – Freiburg<br>37 sub |  | EDSD – Rostock<br>57 sub |  |
| --- | --- | --- | --- | --- |
| Fit | Age | MMSE | Age | MMSE |
| ADNI-2,<br>213 sub | $r=0.49$,<br>**$p=4 \cdot 10^{-3}$**,<br>$p_b=0.05$ | $r=0.16$,<br>$p=0.36$,<br>$p_b=1$ | $r=0.27$,<br>$p=0.05$,<br>$p_b=0.72$ | $r=-0.21$,<br>$p=0.13$,<br>$p_b=1$ |
| OASIS-3,<br>272 sub | $r=0.17$,<br>$p=0.35$,<br>$p_b=1$ | $r=0.45$,<br>**$p=9 \cdot 10^{-3}$**,<br>$p_b=0.12$ | $r=0.18$,<br>$p=0.19$,<br>$p_b=1$ | $r=0.06$,<br>$p=0.64$,<br>$p_b=1$ |
| OASIS-3,<br>139 sub | $r=0.51$,<br>**$p=3 \cdot 10^{-3}$**,<br>**$p_b=0.04$** | $r=0.35$,<br>**$p=0.04$**,<br>$p_b=0.6$ | $r=0.2$,<br>$p=0.15$,<br>$p_b=1$ | $r=0.07$,<br>$p=0.61$,<br>$p_b=1$ |
| OASIS-3,<br>125 sub | $r=0.44$,<br>**$p=0.01$**,<br>$p_b=0.15$ | $r=-0.3$,<br>$p=0.09$,<br>$p_b=1$ | $r=0.22$,<br>$p=0.12$,<br>$p_b=1$ | $r=0.15$,<br>$p=0.27$,<br>$p_b=1$ |
| OASIS-3,<br>116 sub | $r=0.47$,<br>**$p=5 \cdot 10^{-3}$**,<br>$p_b=0.08$ | $r=0.14$,<br>$p=0.45$,<br>$p_b=1$ | $r=0.19$,<br>$p=0.18$,<br>$p_b=1$ | $r=-0.17$,<br>$p=0.24$,<br>$p_b=1$ |
| OASIS-3,<br>All 635 sub | $r=0.25$,<br>$p=0.16$,<br>$p_b=1$ | $r=0.4$,<br>**$p=0.03$**,<br>$p_b=0.35$ | $r=0.19$,<br>$p=0.17$,<br>$p_b=1$ | $r=-0.01$,<br>$p=0.94$,<br>$p_b=1$ |
| HCP,<br>100 sub | $r=0.13$,<br>$p=0.47$,<br>$p_b=1$ | $r=-0.08$,<br>$p=0.64$,<br>$p_b=1$ | $r=0.1$,<br>$p=0.48$,<br>$p_b=1$ | $r=-0.02$,<br>$p=0.89$,<br>$p_b=1$ |

### 3.6. Anti-correlated connectivity results

For ADNI-2, we computed the cross-subject linear correlation coefficient between all pairs of structural connections, keeping *|R^−^|* = 1978 pairs for which *r* := *R_i,j_ < −*0.1. From those, the correlation between the left cortico-subcortical insula-caudate connection and the left corticocortical precentral-entorhinal connection (Figure 6, top, left) was most significant (*p* = 3 *·* 10*^−^*^6^, *p_b_* = 6 *·* 10*^−^*^3^) with *r* = −0.31 and the robust (bisquare) fit slope *m* = −0.40. (The top 20 significant pairs all involved the insula-caudate connection.)

**Figure 6:**
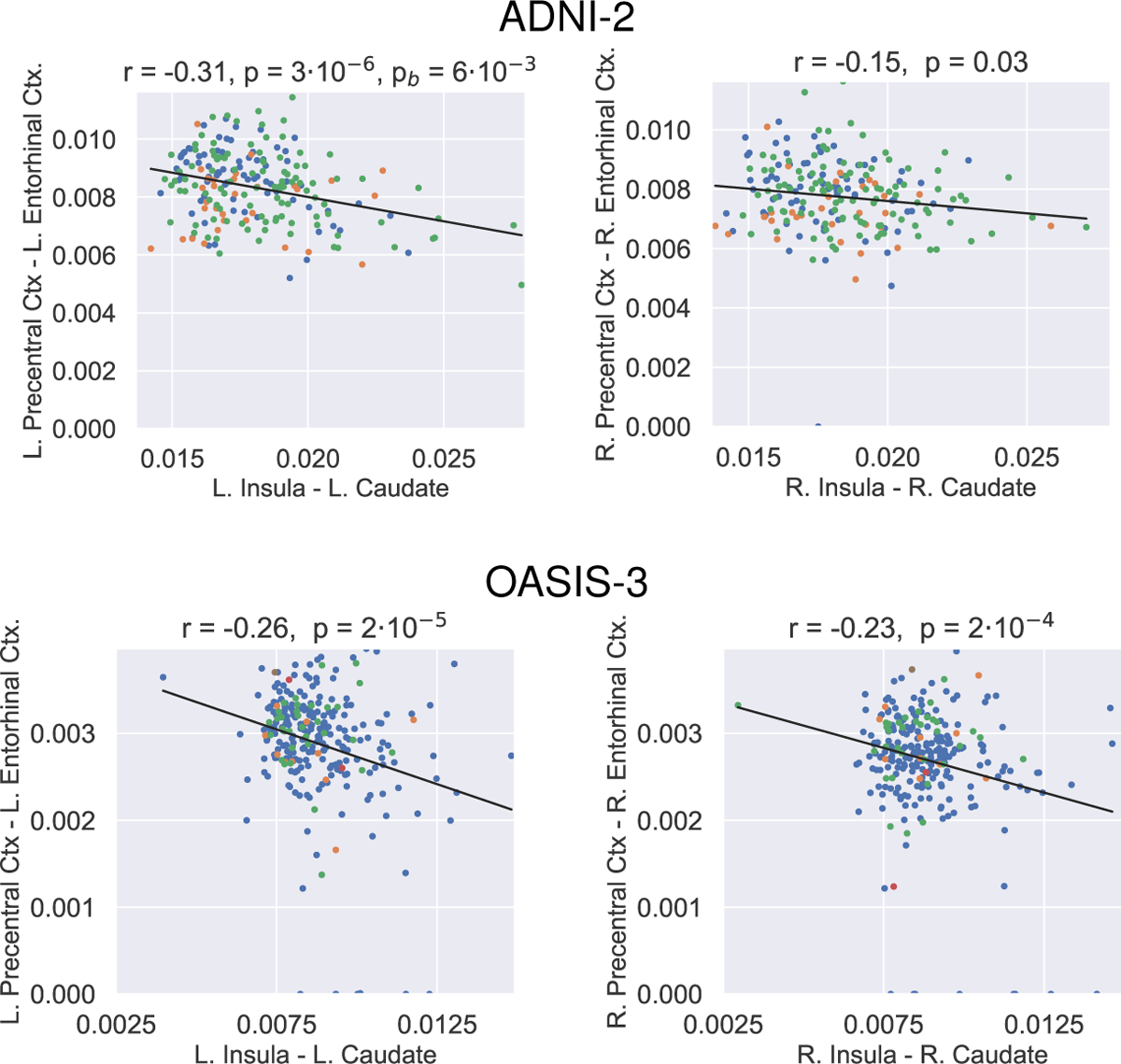
Negative correlation between the insula-caudate and the precentral-entorhinal structural connections in the left and right hemispheres, across ADNI-2 (top) and 272-subject cohort of OASIS-3 (bottom) populations. CDR values are encoded in the colors of the dots: 0 (blue), 0.5 (green), 1 (orange), 2 (red) and 3 (brown) (refer to Figure 3).

We then tested whether the same two connections were inversely correlated also in the right hemisphere, which was true with high significance (*r* = −0.15, *p* = 0.03, *m* = −0.24; Figure 6, top, right). Since here we tested a specific pair of connections in the right hemisphere, correction for multiple comparisons was not necessary.

Next, for external validation and replication, we tested the hypothesis that the pair of insula-caudate and precentral-entorhinal connections are negatively correlated in the first (largest) OASIS-3 database cohort. This hypothesis was validated on this new dataset in both the left (*r* = −0.26, *p* = 2 *·* 10*^−^*^5^, *m* = −0.48) and the right (*r* = −0.23, *p* = 2 *·* 10*^−^*^4^, *m* = −0.28) hemispheres (Figure 6, bottom).

We then computed the correlation of the caudate-insula connection with the CDR and the MMSE score in the OASIS-3 database. While the CDR was negatively correlated with mean connectivity as reported in the previous subsection, it was positively correlated with the caudate-insula connection in the left (*r* = 0.19, *p* = 10*^−^*^3^) and right (*r* = 0.22, *p* = 2 *·* 10*^−^*^4^) hemispheres. Likewise, whereas the MMSE score was positively correlated with mean connectivity, it was negatively correlated with the caudate-insula connection in the left (*r* = −0.12, *p* = 0.046) and right (*r* = −0.12, *p* = 0.04) hemispheres.

### 3.6.1. Null results

In contrast, we did not observe any negative correlation between the insula-caudate and precentral-entorhinal connections across the young-adult HCP subjects^7^. By reversing the order of ADNI-2 and OASIS-3 databases in this experiment, the most significantly anti-correlated pair found in OASIS-3 was not negatively correlated in ADNI-2. In addition, the anti-correlation between the insula-caudate and precentral-entorhinal connections was not observed in OASIS-3 when we included most (652) OASIS-3 subjects, which had heterogeneous scan descriptions (as opposed to the 272-subject cohort).

## 4. Discussion

In this work, we used our previously proposed approach (Frau-Pascual et al., 2019b) to compute and analyze structural brain connectivity in dementia populations. This method models structural connectivity as electric conductance, computing it as a weighted sum of all possible paths between two areas, following the information given by the diffusion tensors. We previously showed (Frau-Pascual et al., 2019b) that this method outperformed deterministic tractography in producing structural connectivity that was more correlated with functional connectivity, possibly due to the fact that all paths – including direct and indirect^8^ – were considered. In this work, we employed our conductance method and investigated its potential in the study of aging and AD. The conductance model was sensitive to AD-related changes in not only diffusion but also geometric properties of the brain WM. For instance, given that this method takes into account distances and paths, changes in subcortical volumes and cortical thickness could also affect the measured connectivity. The conductance might be affected by the GM and WM volume, as they affect ROI sizes and pathways between a pair of ROIs, respectively. Shrinkage in volume could also draw ROIs closer to each other, producing shorter pathways.

Our results were based on the HCP data for healthy young population, and ADNI-2, OASIS-3, and EDSD data for elderly and AD populations. In total, we analyzed 100 young healthy subjects, and 959 elderly subjects, from which 153 had been diagnosed with AD dementia, 122 had MCI, and 83 had other types of dementias or pathologies. This was a heterogeneous pool of subjects with various diagnoses scanned at different sites with different acquisition parameters. Such heterogeneity could make the results more robust and allow for replicability analysis, but also introduce variability that could reduce statistical power.

We used independent variables such as the subject’s age as well as the cognitive scores of CDR and MMSE, which quantify the progression of dementia for a subject, even though they do not clarify whether dementia is due to AD, aging, or other causes. It is worth noting that, as shown in Figure 2, the ways CDR is defined and used in ADNI-2 and OASIS-3 are not identical, and neither are the relationships between CDR and MMSE in these two datasets, as seen when comparing to diagnosis. We did not consider diagnosis when analyzing our data and focused only on these scores (see however Figure 7, left, in the Supplementary Materials for a diagnosis-specific histogram of mean conductance). Previous works have modeled the association between dMRI measures and changes in executive and memory function scores (Scott et al., 2017).

**Figure 7:**
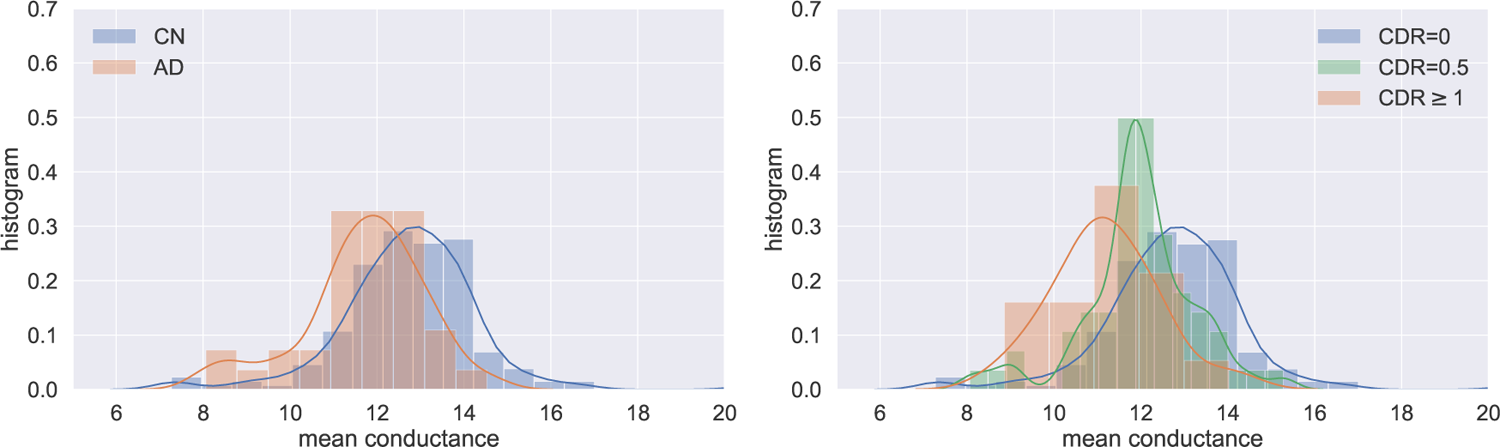
Distribution of mean conductance across disease stage and CDR scales in OASIS-3.

We first summarized the conductance values by averaging them across all region pairs. We considered the independent variables of age, CDR, and MMSE (as well as cortical and subcortical volumes in Figure 8 of the Supplementary Materials). As illustrated in Figure 3, we found expected trends already in the mean conductance: it correlated negatively with age, negatively with CDR, positively with MMSE, and positively with the volumes, except for subcortical regions in ADNI-2. Interestingly, HCP data with healthy young adults followed the same trends in age and volumes. Similar trends have been reported for diffusion measures such as FA and mean diffusivity (Zavaliangos-Petropulu et al., 2019).

**Figure 8:**
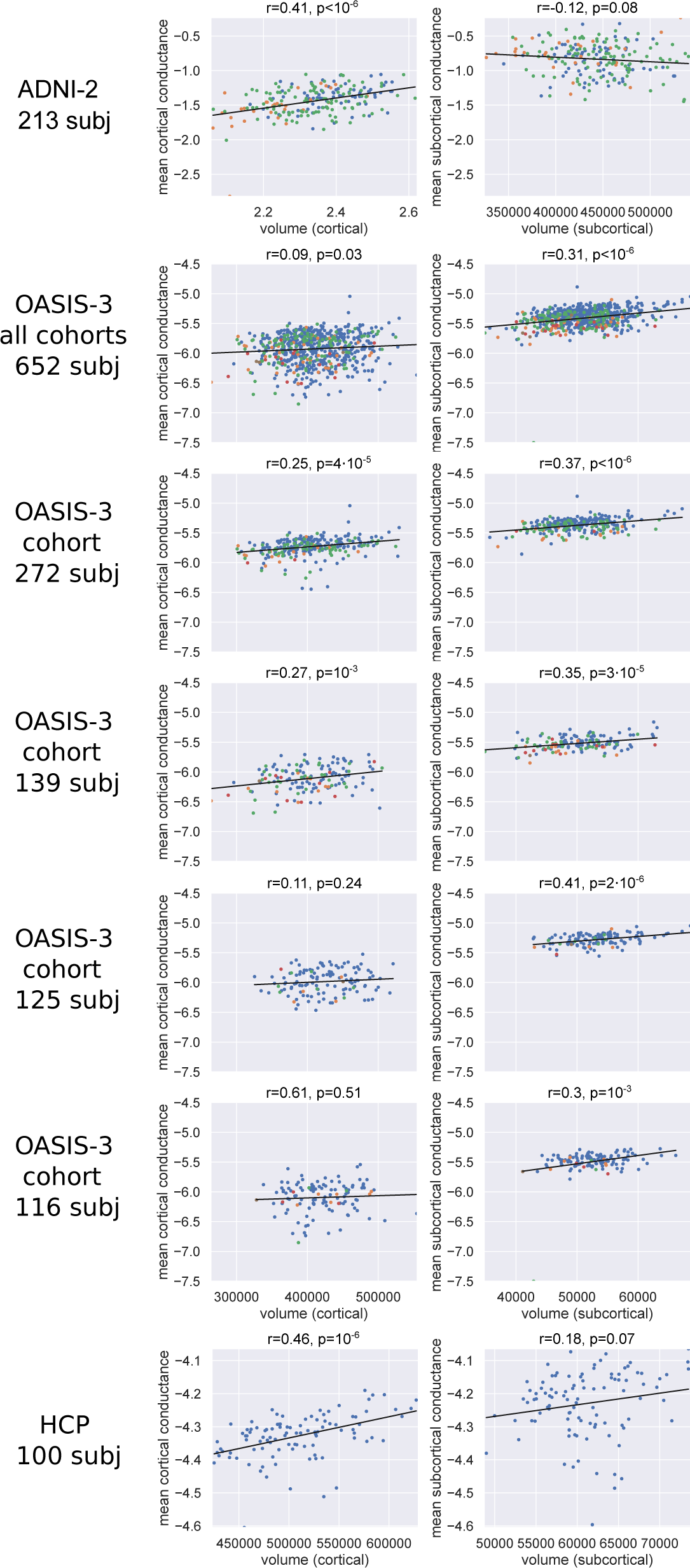
Correlation of mean conductance with cortical and subcortical volumes. Note that the averages were computed only over cortical or subcortical regions.

Next, we investigated the predictive power of mean conductance when training a model and predicting from it, either using the same dataset or different ones. As seen in tables 2 and 3, prediction values were significant for age using most cohorts except for HCP, although in some cases we lost significance after the Bonferroni correction. The generally lower accuracy in cross-prediction compared to within-cohort prediction could be due to difference in scanners and/or inconsistency of imaging protocols used at different imaging sites. Putting all the OASIS-3 data together, in either training or testing, produced significant results. Regarding the HCP, however, most values were not significant, which may have been because the age range of the HCP subjects is not only narrow, but very different from those of the other (dementia) datasets. Predicting cognitive scores of CDR and MMSE from conductance values, as seen in tables 4, 5, 6, and 7, produced significant results mostly only in cohorts with more than 200 subjects. This is probably due to unbalanced values/categories, and, also, the fact that these scores are variable across datasets and diagnoses. When we considered all the OASIS-3 data together, we achieved significant results when training with OASIS-3. Nonetheless, the fact that the prediction was generally significant in datasets with over 200 subjects suggests that sample size may be playing an important role in the prediction power.

We then considered the region-pairwise connections to see how correlated individual connections were with changes in age, and cognitive scores. Previous works have shown correlations of dMRI-derived measures and cognitive scores in the corpus callosum (Moseley et al., 2002), cingulum (Mito et al., 2018), and temporal lobe (Nir et al., 2013). As described in Section 3.4, in OASIS-3 and ADNI-2 the correlation with age was significant for more than a third of the connections, whereas the correlation with cognition was significant in about a fifth of the connections. However, the affected regions were similar in OASIS-3, possibly due to an overlap between cognition and age effects on brain connectivity, and the cognitive decline that accompanies aging. An interesting question here is how much of the correlation of conductance with CDR and MMSE overlaps with the correlation with age and sex. To clarify this nuance, in a different experiment, we regressed out the effects of age and sex and found residual connections that were significantly correlated with MMSE and CDR. In ADNI-2, we observed different patterns of correlation between age and cognitive scores. However, CDR and MMSE correlation patterns did not change when we regressed out the effects of age and sex. It is worth noting that subject head motion has been found to bias results when comparing groups (Yendiki et al., 2014), and that it is likely that in this case it is correlated with cognitive impairment and age.

Once we were done fine-tuning our analysis pipeline, we applied our trained models to two cohorts of the held-out EDSD database for external evaluation. Table 9 shows that the prediction was fairly successful on the Freiburg cohort, but not on the Rostock cohort. We noticed a general trend that the prediction was more accurate among datasets where age and MMSE strongly negatively correlated, such as ADNI-2 (*r*=-0.2, *p*=4*·*10*^−^*^3^), OASIS-3 (*r*=-0.22, *p*=2*·*10*^−^*^8^), and EDSD-Freiburg (*r*=-0.64, *p*=5*·*10*^−^*^5^), than in other datasets, i.e., HCP (*r*=0.04, *p*=0.67) and EDSD-Rostock (*r*=0.14, *p*=0.31). This might mean that for the best prediction results, models must be trained on samples that represent the target population demographically and clinically.

Lastly, we considered the interrelationship among connections.^9^ We found anticorrelation between certain connections, which was consistent across datasets and in both hemispheres. AD patients are known to suffer from connectivity attenuation, but also characterized by brain reorganization and plasticity (Dillen et al., 2016; Kim et al., 2015a). Early in the disease, connectivity within some (i.e., frontal) brain regions increases – possibly due to a compensatory reallocation of cognitive resources – but eventually declines as the disease progresses (Sohn et al., 2014; Brier et al., 2012; Schultz et al., 2017). The connections that we found to be anticorrelated were the precentral-entorhinal cortex and the insula-caudate connections; i.e., the stronger one connection, the weaker the other, across the population. Notably, we observed an increasing trend in the caudate-insula connection strength with respect to CDR. In fact, there is evidence of increased FA in the left caudate in pre-symptomatic familial AD subjects (Ryan et al., 2013), increased structural connectivity in the right insula (Ye et al., 2019), and increased functional connectivity between the frontal lobe and the corpus striatum (Supekar et al., 2008) in AD. In addition, because the conductance method accounts for indirect paths, a possible enhancement in the thalamus and putamen structural connectivity in AD (Ryan et al., 2013; Ye et al., 2019) might also have contributed to the increase in the caudate-insula connectivity. Furthermore, the fact that this negative correlation was observed consistently in the older adults and those on the dementia spectrum (ADNI-2 and OASIS-3), but not in young healthy adults (HCP), suggests that this significant anti-correlation might be due to progression of dementia or aging. Although these results do not necessarily imply a compensatory effect at this stage, our approach may prove useful in a study to discover compensatory connections. Including all OASIS-3 subjects (as opposed to only a subset with homogeneous scans) did not externally validate the anticorrelation hypothesis generated from ADNI-2, possibly because the various acquisition parameters created a large variance in the data that dominated the putative effects. It is important to note that an increase in the measured structural connectivity could stem from factors other than an actual strengthening of the tract. WM atrophy, volume reduction (Ye et al., 2019), and other geometrical variabilities could make ROIs closer to each other, leading to elevated measured structural connectivity. To mitigate this effect, we excluded subcortico-subcortical regions in our study of anti-correlated connections.

Additionally, in regions with fiber crossing, selective axonal loss can increase the FA and subsequently measured structural connectivity (Ryan et al., 2013; Kim et al., 2015b; Douaud et al., 2011).

This article focuses on findings from existing dementia populations using our conductance-based connectivity computation method. For an evaluation of this method, we refer the interested reader to our previous comparisons with existing approaches (Frau-Pascual et al., 2018, 2019b,a; Mohammadi et al., 2020).

## 5. Conclusion

In this work, we focused on the study of the aging and Alzheimer’s disease (AD) populations through connectomics. We applied our conductance method to several databases to detect brain changes related with aging and AD. Results indicated the predictive potential of the conductance measure, especially for age. Although mean conductance values exhibited the expected trends, the prediction of cognitive scores varied across datasets. An important but not surprising finding was that age and cognitive scores of CDR and MMSE largely overlapped. We also correlated brain connections with each other across populations and discovered significantly anti-correlated structural connections. Future work consists of using longitudinal data to further explore the prediction of cognitive scores, and test the hypothesis that anti-correlated connections are indeed compensatory.

## 6. Authorship Confirmation Statement

AF and IA did the project design, writing, and analysis. JA, DV, AY, DS, and BF gave substantial feedback that considerably improved the manuscript.

## Acknowledgements

A preprint of this work can be found at https://www.biorxiv.org/content/10.1101/2020.09.15.298331v4 since 09/15/2020.

## 9. Supplementary Materials

The distribution of mean connectivity for the CN and AD dementia groups, shown in Figure 7, demonstrates an overlap between the two groups, although with separated means (*t* = 4.06*, p* = 8*·*10*^−^*^5^). When we plotted according to the CDR scale, we observed overlapped distributions again, with separated means: *t* = 4.04*, p* = 7*·*10*^−^*^5^ between CDR = 0 and CDR = 0.5; *t* = 2.31*, p* = 7*·*10*^−^*^5^ between CDR = 0.5 and CDR *≥* 1; and *t* = 5.1*, p* = 7 *·* 10*^−^*^6^ between CDR = 0 and CDR *≥* 1. However, classification results were not as significant, probably due to unbalanced cohort size. We are showing only the 272-subject OASIS-3 cohort because the rest of the cohorts did not have enough AD samples.

Table 10 shows the significance of predicting age when using linear regression in only the CN subjects (controls, CDR= 0), in different cohorts. The *p*-values are less significant than when we used all CDR values, probably due to the smaller sample size. When we combined all OASIS-3 data, we got values of *r* = 0.220, *p* = 10*^−^*^6^, *p_b_* = 4 *·* 10*^−^*^6^ for within-cohort prediction, and values of *r* = 0.420, *p* = 10*^−^*^4^, *p_b_* = 9 *·* 10*^−^*^4^ when training on OASIS-3 and testing on ADNI-2, and *r* = 0.175, *p* = 10*^−^*^4^, *p_b_* = 8 *·* 10*^−^*^4^ when training on ADNI-2 and testing on OASIS-3, and a negative *r* when training with OASIS-3 and testing on HCP and vice versa. In this comparison (with all of OASIS-3 in a single cohort), we Bonferroni-corrected (*p_b_*) with a factor 3 in the within-cohort prediction and a factor 6 in the cross-prediction case.

**Table 10:** Prediction of **age** within cohorts in ADNI-2, OASIS-3 (considering only CDR= 0), and HCP.

| ADNI-2 |  | OASIS-3 |  |  | HCP |
| --- | --- | --- | --- | --- | --- |
| 213 sub | 272 sub | 139 sub | 125 sub | 116 sub | 100 sub |
| $r=0.51$,<br>$p=2 \cdot 10^{-6}$,<br>$p_b=10^{-5}$ | $r=0.09$,<br>$p=0.22$,<br>$p_b=1$ | $r=0.47$,<br>$p=10^{-5}$,<br>$p_b=6 \cdot 10^{-5}$ | $r=0.46$,<br>$p=5 \cdot 10^{-7}$,<br>$p_b=3 \cdot 10^{-6}$ | $r=0.35$,<br>$p=5 \cdot 10^{-4}$,<br>$p_b=3 \cdot 10^{-3}$ | $r=-0.06$,<br>$p=0.55$,<br>$p_b=1$ |

**Table 11:** Prediction of **age** with different cohorts in ADNI-2, OASIS-3 (considering only CDR= 0), and HCP. *pb* stands for Bonferroni-corrected *p*-value. *p*-values under 0.05 with a positive *r* are highlighted.

| Fit | Predict |  |  |  |  |  |
| --- | --- | --- | --- | --- | --- | --- |
|  | ADNI-2<br>213 sub | 272 sub | 139 sub | 125 sub | 116 sub | HCP<br>100 sub |
| ADNI-2<br>213 sub | | $r=0.21$,<br>$p=5 \cdot 10^{-3}$,<br>$p_b=0.14$ | $r=0.1$,<br>$p=0.38$,<br>$p_b=1$ | $r=0.09$,<br>$p=0.36$,<br>$p_b=1$ | $r=0.13$,<br>$p=0.2$,<br>$p_b=1$ | $r=-0.11$,<br>$p=0.29$,<br>$p_b=1$ |
| OASIS-3<br>272 sub | $r=0.46$,<br>$p=3 \cdot 10^{-5}$,<br>$p_b=10^{-3}$ | | $r=0.33$,<br>$p=3 \cdot 10^{-3}$,<br>$p_b=0.09$ | $r=0.13$,<br>$p=0.16$,<br>$p_b=1$ | $r=0.29$,<br>$p=4 \cdot 10^{-3}$,<br>$p_b=0.11$ | $r=-0.1$,<br>$p=0.31$,<br>$p_b=1$ |
| OASIS-3<br>139 sub | $r=0.54$,<br>$p=4 \cdot 10^{-7}$,<br>$p_b=10^{-5}$ | $r=0.52$,<br>$p=2 \cdot 10^{-14}$,<br>$p_b=6 \cdot 10^{-13}$ | | $r=0.34$,<br>$p=3 \cdot 10^{-4}$,<br>$p_b=8 \cdot 10^{-3}$ | $r=0.37$,<br>$p=10^{-4}$,<br>$p_b=4 \cdot 10^{-3}$ | $r=0.11$,<br>$p=0.27$,<br>$p_b=1$ |
| OASIS-3<br>125 sub | $r=0.33$,<br>$p=3 \cdot 10^{-3}$,<br>$p_b=0.1$ | $r=0.16$,<br>$p=0.03$,<br>$p_b=0.98$ | $r=0.38$,<br>$p=5 \cdot 10^{-4}$,<br>$p_b=0.01$ | | $r=0.31$,<br>$p=2 \cdot 10^{-3}$,<br>$p_b=0.049$ | $r=-0.0$,<br>$p=0.99$,<br>$p_b=1$ |
| OASIS-3<br>116 sub | $r=0.55$,<br>$p=2 \cdot 10^{-7}$,<br>$p_b=6 \cdot 10^{-6}$ | $r=0.38$,<br>$p=8 \cdot 10^{-8}$,<br>$p_b=2 \cdot 10^{-6}$ | $r=0.55$,<br>$p=9 \cdot 10^{-8}$,<br>$p_b=3 \cdot 10^{-6}$ | $r=0.22$,<br>$p=0.02$,<br>$p_b=0.69$ | | $r=0.27$,<br>$p=6 \cdot 10^{-3}$,<br>$p_b=0.18$ |
| HCP<br>100 sub | $r=-0.38$,<br>$p=6 \cdot 10^{-4}$,<br>$p_b=0.02$ | $r=-0.14$,<br>$p=0.056$,<br>$p_b=1$ | $r=0.03$,<br>$p=0.79$,<br>$p_b=1$ | $r=-0.19$,<br>$p=0.04$,<br>$p_b=1$ | $r=-0.16$,<br>$p=0.11$,<br>$p_b=1$ | |

Figure 8 depicts the correlation of mean conductance with cortical and subcortical volumes, where the averages reported are over only cortical or subcortical conductances, respectively. As in the previous figures, the color-code of the dots reflects the CDR values. The positive *r* in both columns indicate a consistent positive trend of mean cortical conductance and mean subcortical conductance with respect to cortical and subcortical volumes. Only ADNI-2 shows a negative trend for subcortical conductance with respect to volume, although the *p*-value was not significant. Functional connectivity has also been recently correlated with volumes in AD (Sarli et al., 2020). Volumes have been previously used in the prediction of MMSE and CDR decline in MCI (Kovacevic et al., 2009).

## Footnotes

Our codes are publicly available at: www.nitrc.org/projects/conductance

FreeSurfer, https://surfer.nmr.mgh.harvard.edu

FSL, https://fsl.fmrib.ox.ac.uk/fsl/fslwiki

eddy openmp command was used in ADNI-2 and eddy correct in OASIS-3

DSI Studio, http://dsi-studio.labsolver.org

The ADNI (http://adni.loni.usc.edu) was launched in 2003 as a public-private partnership, led by Principal Investigator Michael W. Weiner, MD. The primary goal of ADNI has been to test whether serial MRI, positron emission tomography, other biological markers, and clinical and neuropsychological assessment can be combined to measure the progression of MCI and early AD.

The negative correlation was not observed in functional connectivity either.

One could think of direct and indirect (multi-synaptic) connections as nonstop and multi-stop commercial flights, respectively.

This work has previously been preliminarily presented (Aganj et al., 2020).

